# Structural models predict a significantly higher binding affinity between the NblA protein of cyanophage Ma-LMM01 and the phycocyanin of *Microcystis aeruginosa* NIES-298 compared to the host homolog

**DOI:** 10.1101/2024.07.03.601984

**Authors:** Isaac Meza-Padilla, Brendan J. McConkey, Jozef I. Nissimov

## Abstract

Horizontal gene transfer events between viruses and hosts are widespread across the virosphere. In cyanophage-host systems, such events often involve the transfer of genes involved in photosynthetic processes. The genome of the lytic cyanomyovirus Ma-LMM01 infecting the toxic, bloom-forming, freshwater *Microcystis aeruginosa* NIES-298 contains a homolog of the *non-bleaching A* (*nblA*) gene, which was probably acquired from its host. The function of the NblA protein is to disassemble phycobilisomes, cyanobacterial light harvesting complexes that can comprise up to half of the cellular soluble protein content. NblA thus plays an essential dual role in cyanobacteria: it protects the cell from high light intensities and increases the intracellular nitrogen pool under nutrient limitation. NblA has previously been shown to interact with phycocyanin, one of the main components of phycobilisomes. Using structural modeling and protein-protein docking, we show that the NblA dimer of Ma-LMM01 is predicted to have a significantly higher binding affinity for *M. aeruginosa* NIES-298 phycocyanin (αβ)_6_ hexamers, compared to the host homolog. Protein-protein docking suggests that the viral NblA structural model is able to bind deeper into the phycocyanin groove. The main structural difference between the virus and host NblA appears to be an additional α-helix near the N-terminus of the viral NblA, which could be partly responsible for the deeper binding into phycocyanin. This unique helical region, absent in the cellular NblA, would be expected to constitute a viral evolutionary innovation. We propose that a higher binding affinity of NblA to the host phycocyanin may represent a selective advantage for the virus, whose rapid infection cycle requires an increased phycobilisome degradation rate that is not fulfilled by the NblA of the host.

## 1. Introduction

Aquatic viruses influence global biogeochemical cycles, control the abundance and diversity of their hosts, shape ecological food webs, accelerate coevolutionary processes, and transfer genetic information in marine and freshwater environments (DeLong et al., 2023; Fuhrman, 1999; Rohwer & Thurber, 2009). Virus-host horizontal gene transfer (HGT) events, however, are not unique to aquatic ecosystems. In fact, they take place across the whole virosphere. Exemplary cases include the double jelly-roll major capsid protein of diverse archaeal viruses and bacteriophages within *Varidnaviria*, probably exapted from a family of bacterial enzymes involved in carbohydrate metabolism (Krupovic et al., 2022); the acquisition of cholera toxin genes from the temperate single-stranded DNA bacteriophage CTXΦ by *Vibrio cholerae* (Davis & Waldor, 2003; Waldor & Mekalanos, 1996); as well as several HGTs in eukaryotic viruses, including members of the *Nucleocytoviricota*, and their hosts (Irwin et al., 2022). Collectively, HGTs have played foundational roles in macroevolutionary processes of both viruses and cells throughout the history of life on Earth (Forterre & Prangishvili, 2009; Koonin et al., 2022).

In marine cyanophage-host systems, HGT events often involve the transfer of genes involved in photosynthesis. At the beginning of the 21^st^ century, it was found that S-PM2, a myovirus infecting *Synechococcus* strains, encoded D1 (*psbA*) and D2 (*psbD*) proteins in its genome (Mann et al., 2003; Wilson et al., 1993). D1 and D2 are photosystem II reaction center proteins (Barber, 2013). Lindell et al. (2004) discovered two more myoviruses and a podovirus infecting *Prochlorococcus* that also contained auxiliary metabolic genes encoding for proteins involved in photosynthesis, namely, *psbA*, *psbD*, *hli*, *petE*, and *petF*. *Hli*, *petE*, and *petF* encode high-light-inducible protein, plastocyanin, and ferredoxin, respectively, all components of the photosynthetic process (Gross, 2013; Hanke & Mulo, 2013; Konert et al., 2022). Further, phylogenetic analyses suggested that these genes were not only horizontally transferred from cyanobacterial hosts to viruses, but also that they were probably transferred back to the hosts (Lindell et al., 2004).

Although research on freshwater viruses is relatively limited compared to their marine counterparts, various cases of the presence of photosynthesis-related genes in the genomes of freshwater cyanophages have already been reported (Gao et al., 2012; Meng et al., 2023b; Nadel et al., 2019; Ou et al., 2015; Yoshida et al., 2008). The strain-specific double-stranded DNA lytic myovirus Ma-LMM01 infecting the toxic, bloom-forming *Microcystis aeruginosa* NIES-298 is arguably the most comprehensively characterized freshwater cyanophage to date (e.g., Morimoto et al., 2018; Yoshida et al., 2008; Yoshida et al., 2006). Its genome contains a homolog of the *non-bleaching A* (*nblA*) gene, which was probably horizontally acquired from a *Microcystis* host (Ou et al., 2015; Yoshida et al., 2008). The function of the host NblA protein is to disassemble phycobilisomes, the light harvesting complexes of cyanobacteria, which can comprise up to half of the cellular soluble protein content (Baier et al., 2004; Bienert et al., 2006; Grossman et al., 1993). NblA thus plays an essential dual role in cyanobacteria: it protects the cell from high light intensities and increases the intracellular nitrogen pool under nutrient limitation (Collier & Grossman, 1994; Baier et al., 2004; Grossman et al., 1993). NblA has previously been shown to interact with phycocyanin, one of the main components of phycobilisomes (Bienert et al., 2006; Dines et al., 2008; Karradt et al., 2008; Luque et al., 2003; Nguyen et al., 2017). The fact that the *nblA* gene has been fixed in the Ma-LMM01 population readily suggests that it provides a selective advantage for the virus. Indeed, it is highly transcribed during infection (Honda et al., 2014; Morimoto et al., 2018; Yoshida-Takashima et al., 2012). Here we employ structural modelling and protein-protein docking to investigate which of the two NblAs present in the Ma-LMM01/*M. aeruginosa* NIES-298 virocell (i.e., the virus- or the host-encoded one) is predicted to have a higher binding affinity to the phycocyanin of the host. We elaborate on the potential implications of such a difference in the eco-evolutionary context of this virus-host system.

## 2. Materials & Methods

### 2.1. Structural Bioinformatics Pipeline

The amino acid sequences of Ma-LMM01 NblA (vNblA), *M. aeruginosa* NIES-298 NblA (hNblA), *M. aeruginosa* NIES-298 phycocyanin α-subunit, and *M. aeruginosa* NIES-298 phycocyanin β-subunit were downloaded from the NCBI Protein Database (GenBank accession numbers: BAF36096, GBD54109, GBD54899, and GBD54900, respectively; Sayers et al., 2022). Structural modelling for the vNblA dimer, hNblA dimer, and *M. aeruginosa* NIES-298 phycocyanin (αβ)_6_ hexamer (PC) was performed using AlphaFold2 (AF2) ColabFold (Jumper et al., 2021; Mirdita et al., 2022) with an NVIDIA A100-SXM4-40GB graphics processor.

Secondary structure predictions for vNblA and hNblA were further corroborated using PSIPRED 4.0 (Buchan & Jones, 2019). The sequence alignment between vNblA and hNblA was generated using MUSCLE 3.8 through the EMBL-EBI (Edgar, 2004; Madeira et al., 2022). Entropy-based conservation values were calculated and mapped onto the structural models using AL2CO and University of California, San Francisco (UCSF) ChimeraX (Meng et al., 2023a; Pei & Grishin, 2001). The structural models of the vNblA and hNblA dimers were superimposed using the Matchmaker algorithm implemented in ChimeraX (Meng et al., 2023a) with default parameters, while the pairwise structure comparison tool of the DALI server (Holm et al., 2023) was used to generate single-chain alignments. Hits with a DALI Z-score > 2 (i.e., two standard deviations above expected) were considered significant (Holm & Sander, 1995). Protein-protein docking was carried out using ClusPro 2.0 (Kozakov et al., 2017). To construct the vNblA-PC and hNblA-PC complexes, the PC and NblA structural models were used as the receptor and ligand, respectively. The 30 Balanced models were downloaded and subsequently uploaded to the PRODIGY server (Xue et al., 2016) for binding affinity prediction at 25 °C.

### 2.2. Statistical Analyses

Statistical analyses were conducted using R version 4.3.0 (R Core Team, 2023). In order to investigate whether the vNblA-PC and hNblA-PC predicted complexes had significantly different binding affinities, Welch’s *t*-tests were employed. Prior to analyses, the normality of data was tested using a series of Shapiro-Wilk tests. Gibbs free energy change (ΔG) data, as well as the number of charged-charged (CCs), and polar-nonpolar (PNs) intermolecular contacts at the interface within a threshold distance of 5.5 Å followed normal distributions (*P* > 0.062 in all cases). However, dissociation constant (K_d_) data, and the number of charged-polar, charged-nonpolar, polar-polar, and nonpolar-nonpolar intermolecular contacts (CPs, CNs, PPs, and NNs, respectively) departed from normality. Thus, a log transformation was applied to K_d_, CPs, CNs, PPs, and NNs data. After transformation, Kd, CPs, CNs, PPs, and NNs data followed normal distributions (*P* > 0.063 in all cases). ΔG (kcal mol^−1^), K_d_ (M), CCs, PNs, CPs, CNs, PPs and NNs were used as the dependent variables, and complex (vNblA-PC, hNblA-PC) as the grouping variable. Significance was defined at *P* < 0.05.

## 3. Results

### 3.1. Structural Models

The vNblA and hNblA AF2 structural models can be found in **Fig. 1**. An α-helix, absent in the host model, can be observed near the N-terminus of the viral NblA. Secondary structure prediction also supports the presence of this additional helix in the viral protein, spanning from alanine 3 to glutamic acid 12 (**Fig. 1**h). The structural model of PC, required for protein-protein docking and binding affinity prediction, is shown in **Fig. 1**c,f.

**Fig. 1.**
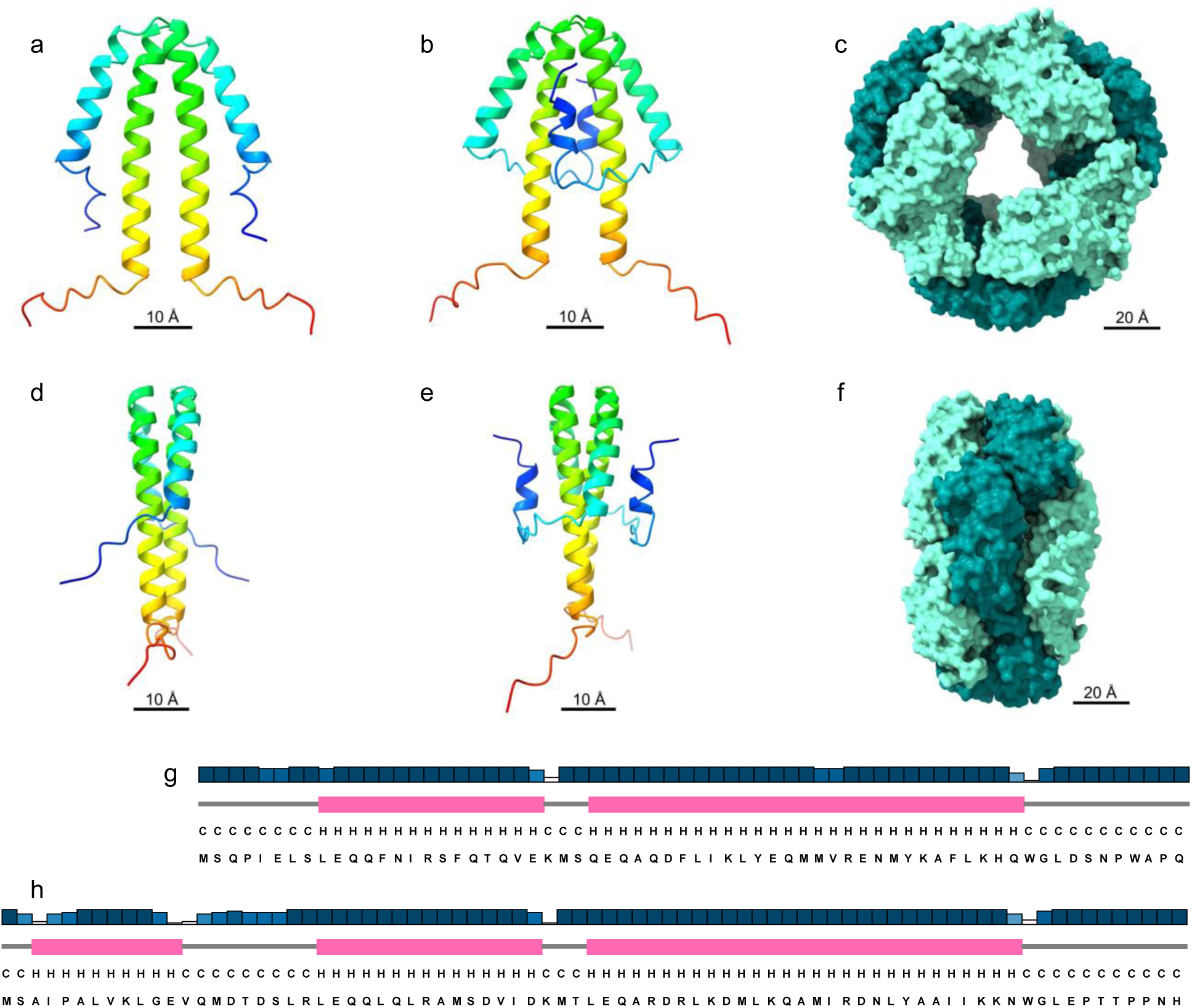
Structural models of the Ma-LMM01/*Microcystis aeruginosa* NIES-298 cyanophage-host system non-bleaching A (NblA) proteins. (a) Front and (d) side views of *M. aeruginosa* NIES-298 NblA (hNblA). (b) Front and (e) side views of Ma-LMM01 NblA (vNblA). NblA models are colored with a rainbow gradient, from the N-terminus in blue to the C-terminus in red. (c) Front and (f) side views of *M. aeruginosa* NIES-298 phycocyanin (αβ)_6_ hexamer (PC). PC α- and β-subunits are colored in dark cyan and aquamarine, respectively. (g) hNblA and (h) vNblA secondary structure predictions. Helices are colored in pink and coils in gray. The confidence of prediction is depicted using a short-tall and white-blue gradient, with short white bars indicating a low prediction confidence, and tall blue bars a high prediction confidence. Note the additional α-helix near the N-terminus of the vNblA secondary and tertiary structural models.

### 3.2. Virus-Host NblA Sequence & Structure Conservation

A moderate pattern of virus-host sequence conservation in the middle region of the vNblA model, where the additional α-helix is also predicted to be located, stands out (**Fig. 2**). These conserved residues and region may be directly involved in the interaction with the host’s phycobilisomes. A significant DALI pairwise structure comparison between vNblA and hNblA (Z-score = 4.4) suggests that the structure of the vNblA model is more conserved compared to its amino acid sequence (**Fig. 2**c,f). Based on the virus-host NblA sequence alignment, the residues absent in hNblA span from leucine 7 to aspartic acid 18, a region containing the predicted α-helix unique to the vNblA model (**Fig. 2**g; section 3.1).

**Fig. 2.**
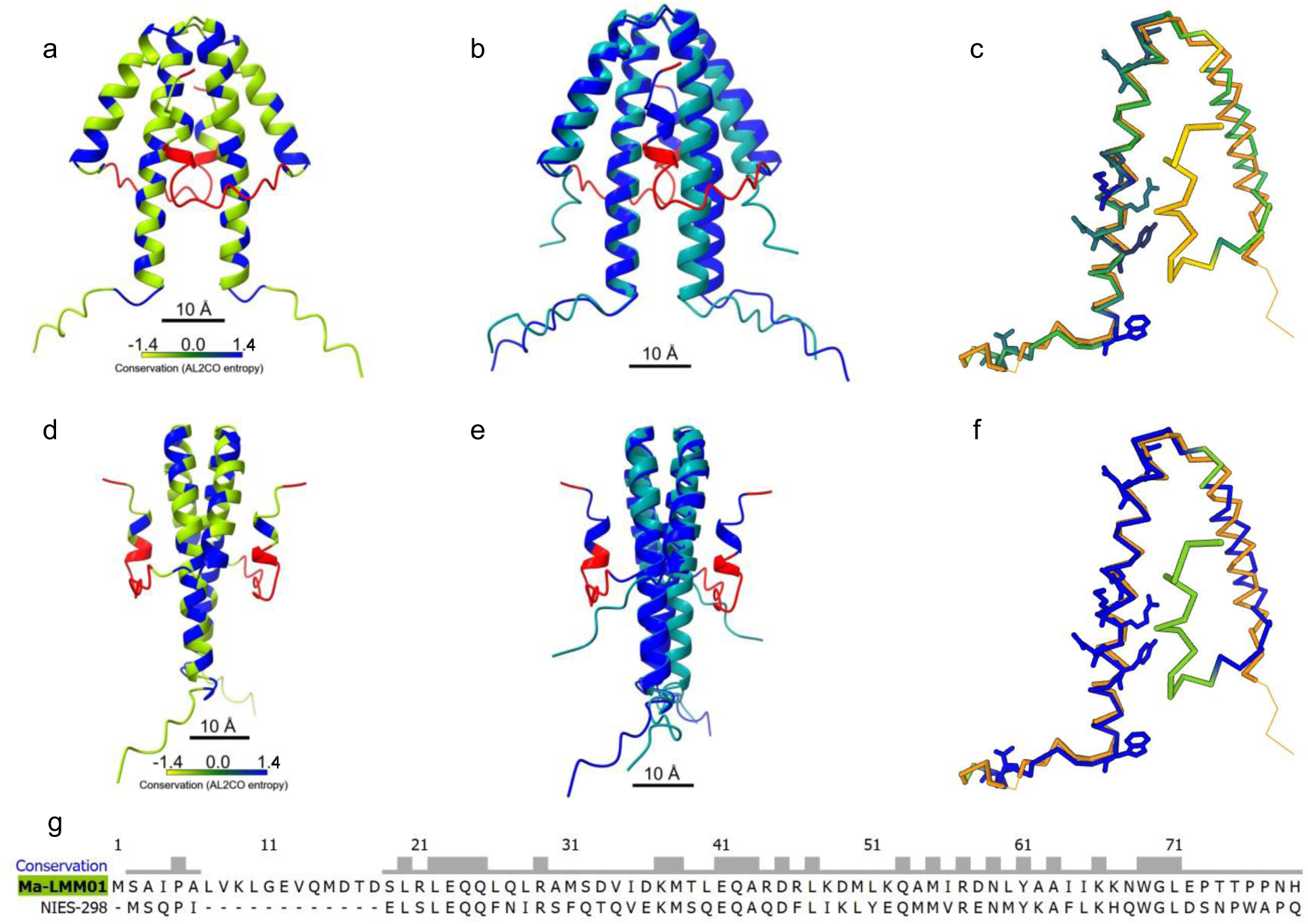
Structure and sequence conservation of the Ma-LMM01/*Microcystis aeruginosa* NIES-298 cyanophage-host system non-bleaching A (NblA) structural models. (a) Front and (d) side views of Ma-LMM01 NblA (vNblA) colored by sequence conservation according to the (g) Ma-LMM01/*M. aeruginosa* NIES-298 NblA sequence alignment. The AL2CO entropy-based amino acid sequence conservation measure between the two proteins is shown using a yellow-green-blue gradient, where yellow and blue represent the least and most conserved residues, respectively. Residues without conservation values, i.e., residues absent in *M. aeruginosa* NIES-298 NblA (hNblA) are shown in red. (b) Front and (e) side views of the vNblA and hNblA superimposed models via the UCSF ChimeraX Matchmaker algorithm. hNblA is shown in dark cyan, while vNblA in blue. As above, the amino acids unique to vNblA are shown in red. Single-chain superimpositions resulting from a DALI pairwise structure comparison with a significant Z-score = 4.4 are displayed in (c) and (f) using a yellow-green-blue scheme mapped onto the C-α trace of the vNblA model, yellow representing the lowest conservation values and blue the highest. In (c) and (f), the relative entropy (0-6.3 bits) was used as the conservation measure. The side chains with a sequence conservation above 4.15 bits are displayed. In (c), vNblA is colored by amino acid sequence conservation; in (f), by structure conservation. hNblA is colored in orange. (g) Ma-LMM01/*M. aeruginosa* NIES-298 NblAs sequence alignment, where the short and tall bars above the residues indicate low and high sequence conservation values, respectively.

### 3.3. Protein-Protein Docking & Binding Affinity Prediction

Statistical analyses indicate that the vNblA-PC and hNblA-PC predicted complexes have significantly different binding affinities, as determined by ΔG (Welch’s *t*-test, *t*_49,667_ = 3.017, *P* = 0.004; **Fig. 3**a) and K_d_ (*t*_49,578_ = 3.038, *P* = 0.004; **Fig. 3**b). Likewise, the predicted complexes have significantly different CC, CP, CN and PP intermolecular contacts at the interface within a threshold distance of 5.5 Å (*P* < 0.041 in all cases; **Fig. 3**c-f). No significant differences were detected regarding the number of PN and NN intermolecular contacts (*P* > 0.05 in both cases). This suggests that the lower ΔG and K_d_ values of the vNblA-PC complex are mainly due to an increased number of CC, CP, and CN intermolecular contacts. ΔG was found to be −15.89 ± 2.69 kcal mol^−1^ (*n* = 30) for the hNblA-PC complex, and −17.66 ± 1.74 kcal mol^−1^ (*n* = 30) for the vNblA-PC complex; while K_d_ was 2.32 × 10^−10^ ± 5.80 × 10^−10^ M (*n* = 30) for the hNblA-PC complex, and 6.86 × 10^−12^ ± 3.28 × 10^−11^ M (*n* = 30) for the vNblA-PC complex. Both of these significantly lower ΔG and K_d_ values of the vNblA-PC predicted complex translate into a significantly higher binding affinity between the vNblA and PC structural models compared to the hNblA and PC models.

**Fig. 3.**
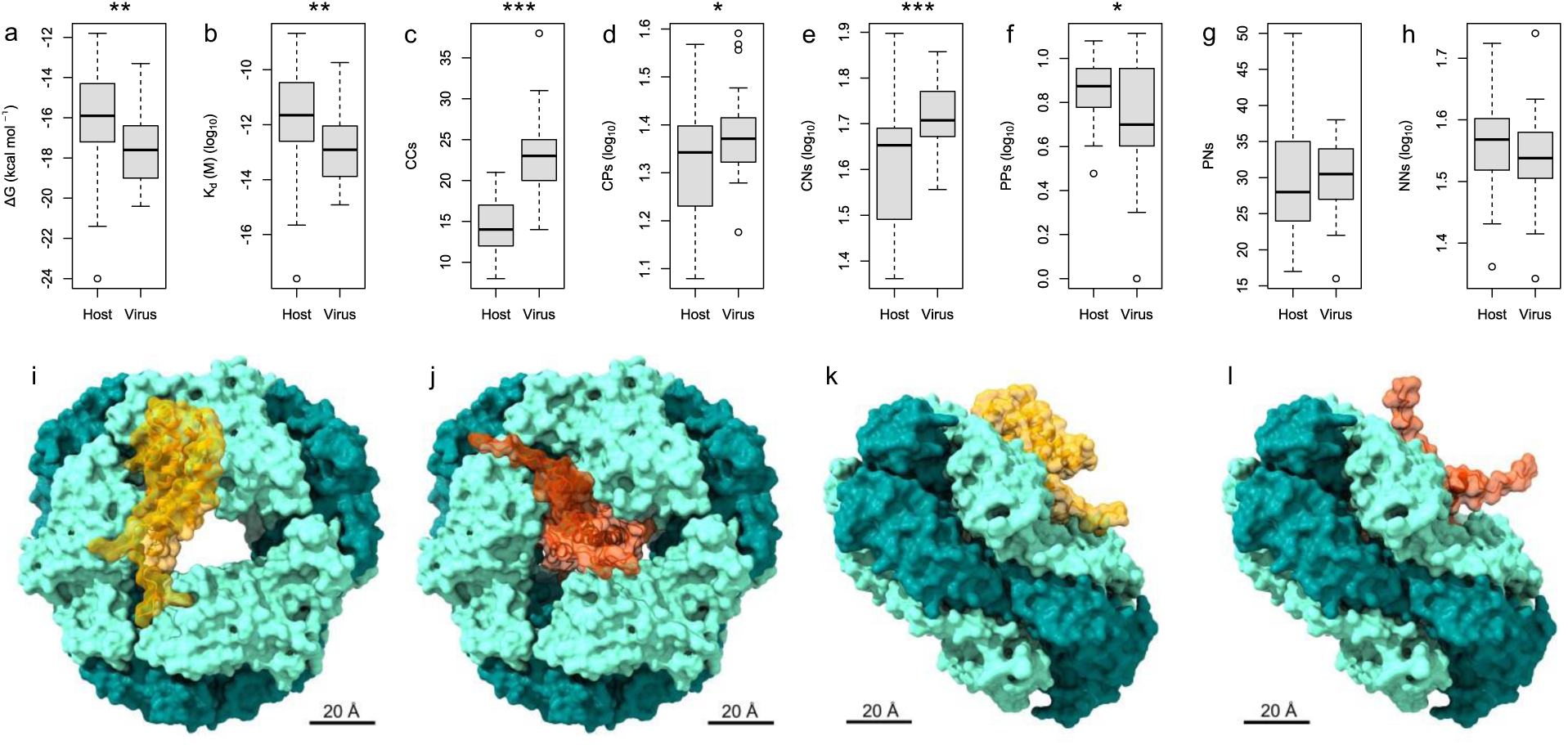
Protein-protein docking and binding affinity prediction between Ma-LMM01 non-bleaching A dimer (vNblA) and *Microcystis aeruginosa* NIES-298 phycocyanin (αβ)_6_ hexamer (PC) structural models compared to *M. aeruginosa* NIES-298 NblA dimer (hNblA) and PC models. (a) ΔG (kcal mol^−1^) and (b) K_d_ (M) of hNblA-PC (host) and vNblA-PC (virus) predicted complexes. (c) Number of host and virus charged-charged (CC), (d) charged-polar (CP), (e) charged-nonpolar (CN), (f) polar-polar (PP), (g) polar-nonpolar (PN), and (h) nonpolar-nonpolar (NN) intermolecular contacts at the interface within a threshold distance of 5.5 Å. See text for statistical values. (i) Front and (k) side views of the best ranked ClusPro 2.0 hNblA-PC docking model. (j) Front and (l) side views of the best ranked vNblA-PC model. hNblA is colored in orange, while vNblA in orange red; PC α- and β-subunits are colored in dark cyan and aquamarine, respectively. Note how vNblA is predicted to bind deeper into the PC groove compared to hNblA. See text for binding affinities of the displayed docking models.

The best ranked hNblA-PC ClusPro 2.0 docking model (**Fig. 3**i,j) was predicted to have ΔG and K_d_ values of −14.9 kcal mol^−1^ and 1.20 × 10^−11^ M, respectively. In comparison, the best ranked vNblA-PC model had a ΔG of −16.3 kcal mol^−1^ and a K_d_ of 1.10 × 10^−12^ M, almost an order of magnitude lower than the hNblA-PC model. It is worth noting that in most complexes the viral and cellular NblAs were in similar positions as they were in the best ranked models (**Fig. S1**), showing consistency in the docking predictions. Most hNblA models (23/30) were bound superficially (compared to vNblA) to PC, with a smaller region inside the PC groove. In contrast, virtually all vNblA models were bound deep inside the PC groove (29/30). The main structural difference between the two NblA models, namely, the additional vNblA α-helix, may contribute, at least in part, to binding vNblA deeper into the PC groove.

## 4. Discussion

The sequencing of the Ma-LMM01 genome (Yoshida et al., 2008) and more recently its *M. aeruginosa* NIES-298 host (Yamaguchi et al., 2018), along with the development of highly accurate structure prediction (Jumper et al., 2021) and other robust structural bioinformatics tools (Kozakov et al., 2017; Xue et al., 2016) provide a unique opportunity to model the NblA and PC structures of this cyanophage-host system and compare their predicted binding affinities. First, a qualitative analysis of the viral and host NblA models readily highlights an additional α-helix present in the viral structural model (section 3.1; **Fig. 1**). Phylogenetic evidence suggests that Ma-LMM01 *nblA* was horizontally acquired from a *Microcystis* host (Ou et al., 2015). Structure conservation analyses, including a significant DALI pairwise structure comparison between vNblA and hNblA (section 3.2; **Fig. 2**), support virus-host structural homology as well. Hence, either the viral *nblA* expanded (insertions) or the host *nblA* shortened (deletions) after the HGT took place. Multiple sequence alignments show that, unlike Ma-LMM01 NblA, cellular NblA amino acid sequences have a similar length (Ou et al., 2015; Yoshida et al., 2008). This arguably renders the expansion of Ma-LMM01 *nblA* more parsimonious compared to the reduction of different cellular *nblA* genes. It follows then that not only has the *nblA* gene become fixed in the Ma-LMM01 population, but it has also expanded throughout evolutionary timescales; the additional vNblA α-helix constitutes a viral evolutionary innovation. Here it may be possible to retrospectively appreciate two steps of the origin of evolutionary novelties (Blount et al., 2012): actualization, represented by the *nblA* HGT event itself; and refinement, represented by the expansion of the viral *nblA* gene.

An increase in genome size (the acquisition of the cellular *nblA* gene by the virus and, to a lesser extent, its expansion) must constitute a selective advantage strong enough to counter the possible trade-off associated with the greater amount of time and resources required to replicate the viral genome (DiMaio, 2012; Edwards et al., 2021; Mills et al., 1967). We propose that such an advantage is the increased binding affinity that the viral NblA is predicted to have for PC compared to the host’s NblA (see section 3.3). The quantitative analyses conducted on the vNblA-PC and hNblA-PC predicted complexes indicate a significantly lower ΔG and K_d_ for the vNblA-PC complex, which translate into a higher binding affinity. Not surprisingly, the number of intermolecular contacts was also predicted to be greater in vNblA-PC compared to the hNblA-PC complex in most contact property categories. The exact binding mechanism of NblA in phycobilisomes has not been resolved yet. However, various studies have provided insights into the mode of action of NblA (Baier et al., 2004; Bienert et al., 2006; Dines et al., 2008; Hu et al., 2020; Karradt et al., 2008; Levi et al., 2018; Luque et al., 2003; Nguyen et al., 2017; Sendersky et al., 2015; Sendersky et al., 2014). For example, Bienert et al. (2006) determined the crystal structure of NblA from *Anabaena* sp. PCC 7120, and using pull-down assays, found that NblA bound the α-subunits of phycocyanin and phycoerythrocyanin; while Nguyen et al. (2017) detected a protein interaction between *S. elongatus* UTEX 2973 NblA and β-phycocyanin. Dines et al. (2008) determined the NblA structures of *S. elongatus* PCC 7942 and

*Thermosynechococcus vulcanus*, and using random mutagenesis, found that amino acids essential for the interaction with phycobilisome proteins were located across the two NblA helices. The hNblA-PC docking models from the present study are consistent with those reported by Nguyen et al. (2017), where *S. elongatus* PCC 7942 NblA and PC X-ray structures docked similarly to *M. aeruginosa* NIES-298 NblA and PC structural models. In contrast, the vNblA-PC docking models differ drastically from both hNblA-PC and those reported previously for *S. elongatus* (Nguyen et al., 2017). In particular, the vNblA model was predicted to bind deeper into the PC groove. The additional α-helix, which appears to be the main structural difference between vNblA and the cellular NblAs, may be partly responsible for this.

Beyond the specific binding site of the viral NblA protein, a higher binding affinity for PC would be expected to increase the disassembling rate of the host phycobilisomes. This could be essential for the virus if the performance of the host protein is not enough during infection. In fact, the viral *nblA* is highly transcribed in the Ma-LMM01/*M. aeruginosa* NIES-298 virocell (Honda et al., 2014; Morimoto et al., 2018; Yoshida-Takashima et al., 2012). The infection cycle of Ma-LMM01 is rapid; the latent period is estimated to take between six to twelve hours (Yoshida et al., 2006). As previously described, phycobilisomes are the light harvesting complexes of cyanobacteria, and can comprise up to half of the cellular soluble protein content (Baier et al., 2004; Bienert et al., 2006; Grossman et al., 1993). A higher phycobilisome degradation rate in the Ma-LMM01/*M. aeruginosa* NIES-298 virocell could lead to a larger burst size after a steeper increase in the intracellular nitrogen pool (Morimoto et al., 2020; Morimoto et al., 2018; Yoshida et al., 2008). A faster degradation of phycobilisomes would also imply a more efficient protection for the virocell from high light intensities to which *M. aeruginosa* NIES-298 is highly susceptible (Morimoto et al., 2020; Morimoto et al., 2018; Yoshida et al., 2008).

Based on the significantly lower ΔG and K_d_ of the vNblA-PC docking models, we conclude that vNblA is predicted to have a higher binding affinity for PC compared to hNblA. The potential implications of this are in line with the biology of Ma-LMM01. However, rather than aiming to provide a definitive answer regarding the mode of action of the vNblA protein, these structural models invite further investigation through experimental approaches, including structure determination efforts. They also highlight the statistics-based hypothesis generating potential of the structural bioinformatics pipeline implemented here, particularly for comparative interactomics in the fields of evolutionary virology and viral ecology.

## Acknowledgements

This study would not have been possible without the continuous support of Dr. Medardo Meza Olea and Dr. Esther Padilla Calderón.

## Data availability

The binding affinity prediction data is available in Supplementary Material (**Tables S1**, **S2**). The structural models can be generated following the structural bioinformatics pipeline described in section 2.1. Alternatively, the models can be provided upon reasonable request to the corresponding author.

## Author contributions

Meza-Padilla conceived the idea, carried out the structural bioinformatics pipeline, conducted and reported the statistical analyses, analyzed the data, and wrote the manuscript. McConkey provided additional insights on structural modeling and molecular docking. Nissimov administered the project, provided resources and funding, and supervised the work. All authors contributed to the refinement (review and editing) of the final manuscript prior to its submission and publication.

## References

Baier, K., Lehmann, H., Stephan, D. P., & Lockau, W. (2004). NblA is essential for phycobilisome degradation in *Anabaena* sp. strain PCC 7120 but not for development of functional heterocysts. Microbiology, 150(8), 2739–2749. 10.1099/mic.0.27153-0.

Barber, J. (2013). Photosystem II: Redox and protein components. In W. J. Lennarz, & M. D. Lane (Eds.), Encyclopedia of Biological Chemistry II. Elsevier. 10.1016/B978-0-12-378630-2.00381-9.

Bienert, R., Baier, K., Volkmer, R., Lockau, W., & Heinemann, U. (2006). Crystal structure of NblA from *Anabaena* sp. PCC 7120, a small protein playing a key role in phycobilisome degradation. Journal of Biological Chemistry, 281(8), 5216–5223. 10.1074/jbc.M507243200.

Blount, Z. D., Barrick, J. E., Davidson, C. J., & Lenski, R. E. (2012). Genomic analysis of a key innovation in an experimental *Escherichia coli* population. Nature, 489(7417), 513–518. 10.1038/nature11514.

Buchan, D. W., & Jones, D. T. (2019). The PSIPRED protein analysis workbench: 20 years on. Nucleic Acids Research, 47(W1), W402–W407. 10.1093/nar/gkz297.

Collier, J. L., & Grossman, A. (1994). A small polypeptide triggers complete degradation of light-harvesting phycobiliproteins in nutrient-deprived cyanobacteria. The EMBO Journal, 13(5), 1039–1047. 10.1002/j.1460-2075.1994.tb06352.x.

Davis, B. M., & Waldor, M. K. (2003). Filamentous phages linked to virulence of *Vibrio cholerae*. Current Opinion in Microbiology, 6(1), 35–42. 10.1016/S1369-5274(02)00005-X.

DeLong, J. P., Van Etten, J. L., & Dunigan, D. D. (2023). Lessons from chloroviruses: the complex and diverse roles of viruses in food webs. Journal of Virology, 97(5), e00275–23. 10.1128/jvi.00275-23.

DiMaio, D. (2012). Viruses, masters at downsizing. Cell Host & Microbe, 11(6), 560–561. 10.1016/j.chom.2012.05.004.

Dines, M., Sendersky, E., David, L., Schwarz, R., & Adir, N. (2008). Structural, functional, and mutational analysis of the NblA protein provides insight into possible modes of interaction with the phycobilisome. Journal of Biological Chemistry, 283(44), 30330–30340. 10.1074/jbc.M804241200.

Edgar, R. C. (2004). MUSCLE: a multiple sequence alignment method with reduced time and space complexity. BMC Bioinformatics, 5, 1–19. 10.1186/1471-2105-5-113.

Edwards, K. F., Steward, G. F., & Schvarcz, C. R. (2021). Making sense of virus size and the tradeoffs shaping viral fitness. Ecology Letters, 24(2), 363–373. 10.1111/ele.13630.

Forterre, P., & Prangishvili, D. (2009). The great billion-year war between ribosome-and capsid-encoding organisms (cells and viruses) as the major source of evolutionary novelties. Annals of the New York Academy of Sciences, 1178(1), 65–77. 10.1111/j.1749-6632.2009.04993.x.

Fuhrman J. A. (1999). Marine viruses and their biogeochemical and ecological effects. Nature, 399(6736), 541–548. 10.1038/21119.

Gao, E. B., Gui, J. F., & Zhang, Q. Y. (2012). A novel cyanophage with a cyanobacterial nonbleaching protein A gene in the genome. Journal of Virology, 86(1), 236–245. 10.1128/jvi.06282-11.

Gross, E. L. (2013). Plastocyanin. In W. J. Lennarz, & M. D. Lane (Eds.), Encyclopedia of Biological Chemistry II. Elsevier. 10.1016/B978-0-12-378630-2.00137-7.

Grossman, A. R., Schaefer, M. R., Chiang, G. G., & Collier, J. (1993). The phycobilisome, a light-harvesting complex responsive to environmental conditions. Microbiological Reviews, 57(3), 725–749. 10.1128/mr.57.3.725-749.1993.

Hanke, G. U. Y., & Mulo, P. (2013). Plant type ferredoxins and ferredoxin-dependent metabolism. Plant, Cell & Environment, 36(6), 1071–1084. 10.1111/pce.12046.

Holm, L., Laiho, A., Törönen, P., & Salgado, M. (2023). DALI shines a light on remote homologs: One hundred discoveries. Protein Science, 32(1), e4519. 10.1002/pro.4519.

Holm, L., & Sander, C. (1995). Dali: a network tool for protein structure comparison. Trends in Biochemical Sciences, 20(11), 478–480. 10.1016/S0968-0004(00)89105-7.

Honda, T., Takahashi, H., Sako, Y., & Yoshida, T. (2014). Gene expression of *Microcystis aeruginosa* during infection of cyanomyovirus Ma-LMM01. Fisheries Science, 80, 83–91.

Hu, P. P., Hou, J. Y., Xu, Y. L., Niu, N. N., Zhao, C., Lu, L., Zhou, M., Scheer, H., & Zhao, K. H. (2020). The role of lyases, NblA and NblB proteins and bilin chromophore transfer in restructuring the cyanobacterial light-harvesting complex. The Plant Journal, 102(3), 529–540. 10.1111/tpj.14647.

Irwin, N. A., Pittis, A. A., Richards, T. A., & Keeling, P. J. (2022). Systematic evaluation of horizontal gene transfer between eukaryotes and viruses. Nature Microbiology, 7(2), 327–336. 10.1038/s41564-021-01026-3.

Jumper, J., Evans, R., Pritzel, A., Green, T., Figurnov, M., Ronneberger, O., Tunyasuvunakool K., Bates, R., Žídek, A., Potapenko, A., Bridgland, A., Meyer, C., Kohl, A. A. S., Ballard, A. J., Cowie, A., Romera-Paredes, B., Nikolov, S., Jain, R., Adler, J., Back, T., Petersen, S., Reiman, D., Clancy, E., Zielinski M., Steinegger, M., Pacholska, M., Berghammer, T., Bodenstein, S., Silver, D., Vinyals, O., Senior, A. W., Kavukcuoglu, K., Kohli, P., & Hassabis, D. (2021). Highly accurate protein structure prediction with AlphaFold. Nature, 596(7873), 583–589. 10.1038/s41586-021-03819-2.

Karradt, A., Sobanski, J., Mattow, J., Lockau, W., & Baier, K. (2008). NblA, a key protein of phycobilisome degradation, interacts with ClpC, a HSP100 chaperone partner of a cyanobacterial Clp protease. Journal of Biological Chemistry, 283(47), 32394–32403. 10.1074/jbc.M805823200.

Konert, M. M., Wysocka, A., Koník, P., & Sobotka, R. (2022). High-light-inducible proteins HliA and HliB: pigment binding and protein–protein interactions. Photosynthesis Research, 152(3), 317–332. 10.1007/s11120-022-00904-z.

Koonin, E. V., Dolja, V. V., & Krupovic, M. (2022). The logic of virus evolution. Cell Host & Microbe, 30(7), 917–929. 10.1016/j.chom.2022.06.008.

Kozakov, D., Hall, D. R., Xia, B., Porter, K. A., Padhorny, D., Yueh, C., Beglov, D., & Vajda, S. (2017). The ClusPro web server for protein–protein docking. Nature Protocols, 12(2), 255–278. 10.1038/nprot.2016.169.

Krupovic, M., Makarova, K. S., & Koonin, E. V. (2022). Cellular homologs of the double jelly-roll major capsid proteins clarify the origins of an ancient virus kingdom. Proceedings of the National Academy of Sciences, 119(5), e2120620119. 10.1073/pnas.2120620119.

Levi, M., Sendersky, E., & Schwarz, R. (2018). Decomposition of cyanobacterial light harvesting complexes: NblA-dependent role of the bilin lyase homolog NblB. The Plant Journal, 94(5), 813–821. 10.1111/tpj.13896.

Lindell, D., Sullivan, M. B., Johnson, Z. I., Tolonen, A. C., Rohwer, F., & Chisholm, S. W. (2004). Transfer of photosynthesis genes to and from *Prochlorococcus* viruses. Proceedings of the National Academy of Sciences, 101(30), 11013–11018. 10.1073/pnas.0401526101.

Luque, I., Ochoa de Alda, J. A., Richaud, C., Zabulon, G., Thomas, J. C., & Houmard, J. (2003). The NblAI protein from the filamentous cyanobacterium *Tolypothrix* PCC 7601: regulation of its expression and interactions with phycobilisome components. Molecular Microbiology, 50(3), 1043–1054. 10.1046/j.1365-2958.2003.03768.x.

Madeira, F., Pearce, M., Tivey, A. R., Basutkar, P., Lee, J., Edbali, O., Madhusoodanan, N., Kolesnikov, A., & Lopez, R. (2022). Search and sequence analysis tools services from EMBL-EBI in 2022. Nucleic Acids Research, 50(W1), W276–W279. 10.1093/nar/gkac240.

Meng, E. C., Goddard, T. D., Pettersen, E. F., Couch, G. S., Pearson, Z. J., Morris, J. H., & Ferrin, T. E. (2023a). UCSF ChimeraX: Tools for structure building and analysis. Protein Science, 32(11), e4792. 10.1002/pro.4792.

Meng, L. H., Ke, F., Zhang, Q. Y., & Zhao, Z. (2023b). Biological and genomic characteristics of MaMV-DH01, a novel freshwater *Myoviridae* cyanophage strain. Microbiology Spectrum, 11(1), e02888–22. 10.1128/spectrum.02888-22.

Mills, D. R., Peterson, R. L., & Spiegelman, S. (1967). An extracellular Darwinian experiment with a self-duplicating nucleic acid molecule. Proceedings of the National Academy of Sciences, 58(1), 217–224. 10.1073/pnas.58.1.217.

Mirdita, M., Schütze, K., Moriwaki, Y., Heo, L., Ovchinnikov, S., & Steinegger, M. (2022). ColabFold: making protein folding accessible to all. Nature Methods, 19(6), 679–682. 10.1038/s41592-022-01488-1.

Morimoto, D., Kimura, S., Sako, Y., & Yoshida, T. (2018). Transcriptome analysis of a bloom-forming cyanobacterium *Microcystis aeruginosa* during Ma-LMM01 phage infection. Frontiers in Microbiology, 9, 317806. 10.3389/fmicb.2018.00002.

Morimoto, D., Šulčius, S., & Yoshida, T. (2020). Viruses of freshwater bloom-forming cyanobacteria: genomic features, infection strategies and coexistence with the host. Environmental Microbiology Reports, 12(5), 486–502. 10.1111/1758-2229.12872o.

Nadel, O., Rozenberg, A., Flores-Uribe, J., Larom, S., Schwarz, R., & Béjà, O. (2019). An uncultured marine cyanophage encodes an active phycobilisome proteolysis adaptor protein NblA. Environmental Microbiology Reports, 11(6), 848–854. 10.1111/1758-2229.12798.

Nguyen, A. Y., Bricker, W. P., Zhang, H., Weisz, D. A., Gross, M. L., & Pakrasi, H. B. (2017). The proteolysis adaptor, NblA, binds to the N-terminus of β-phycocyanin: Implications for the mechanism of phycobilisome degradation. Photosynthesis Research, 132, 95–106. 10.1007/s11120-016-0334-y.

Ou, T., Gao, X. C., Li, S. H., & Zhang, Q. Y. (2015). Genome analysis and gene *nblA* identification of *Microcystis aeruginosa* myovirus (MaMV-DC) reveal the evidence for horizontal gene transfer events between cyanomyovirus and host. Journal of General Virology, 96(12), 3681–3697. 10.1099/jgv.0.000290.

Pei, J., & Grishin, N. V. (2001). AL2CO: calculation of positional conservation in a protein sequence alignment. Bioinformatics, 17(8), 700–712. 10.1093/bioinformatics/17.8.700.

R Core Team. (2023). R: A Language and Environment for Statistical Computing. R Foundation for Statistical Computing, Vienna, Austria. https://www.R-project.org/.

Rohwer, F., & Thurber, R. V. (2009). Viruses manipulate the marine environment. Nature, 459(7244), 207–212. 10.1038/nature08060.

Sayers, E. W., Beck, J., Bolton, E. E., Bourexis, D., Brister, J. R., Canese, K., Chan, J., Comeau, D. C., Connor, R., Funk, K., Kelly, C., Kim, S., Madej T., Marchler-Bauer, A., Lanczycki, C., Lathrop, S., Lu, Z., Thibaud-Nissen, F., Murphy, T., Phan, L., Skripchenko, Y., Tse, T., Wang, J., Williams, R., Trawick, B., W., Pruitt, K., D., & Sherry, S. T. (2022). Database resources of the national center for biotechnology information. Nucleic Acids Research, 50(D1), D20–D26. 10.1093/nar/gkab1112.

Sendersky, E., Kozer, N., Levi, M., Garini, Y., Shav-Tal, Y., & Schwarz, R. (2014). The proteolysis adaptor, NblA, initiates protein pigment degradation by interacting with the cyanobacterial light-harvesting complexes. The Plant Journal, 79(1), 118–126. 10.1111/tpj.12543.

Sendersky, E., Kozer, N., Levi, M., Moizik, M., Garini, Y., Shav-Tal, Y., & Schwarz, R. (2015). The proteolysis adaptor, NblA, is essential for degradation of the core pigment of the cyanobacterial light-harvesting complex. The Plant Journal, 83(5), 845–852. 10.1111/tpj.12931.

Waldor, M. K., & Mekalanos, J. J. (1996). Lysogenic conversion by a filamentous phage encoding cholera toxin. Science, 272(5270), 1910–1914. 10.1126/science.272.5270.1910.

Xue, L. C., Rodrigues, J. P., Kastritis, P. L., Bonvin, A. M., & Vangone, A. (2016). PRODIGY: a web server for predicting the binding affinity of protein–protein complexes. Bioinformatics, 32(23), 3676–3678. 10.1093/bioinformatics/btw514.

Yamaguchi, H., Suzuki, S., & Kawachi, M. (2018). Improved draft genome sequence of *Microcystis aeruginosa* NIES-298, a microcystin-producing cyanobacterium from Lake Kasumigaura, Japan. Genome Announcements, 6(5), 10–1128. 10.1128/genomea.01551-17.

Yoshida, T., Nagasaki, K., Takashima, Y., Shirai, Y., Tomaru, Y., Takao, Y., Sakamoto, S., Hiroishi, S., & Ogata, H. (2008). Ma-LMM01 infecting toxic *Microcystis aeruginosa* illuminates diverse cyanophage genome strategies. Journal of Bacteriology, 190(5), 1762–1772. 10.1128/jb.01534-07.

Yoshida-Takashima, Y., Yoshida, M., Ogata, H., Nagasaki, K., Hiroishi, S., & Yoshida, T. (2012). Cyanophage infection in the bloom-forming cyanobacteria *Microcystis aeruginosa* in surface freshwater. Microbes and Environments, 27(4), 350–355. 10.1264/jsme2.ME12037.

Yoshida, T., Takashima, Y., Tomaru, Y., Shirai, Y., Takao, Y., Hiroishi, S., & Nagasaki, K. (2006). Isolation and characterization of a cyanophage infecting the toxic cyanobacterium *Microcystis aeruginosa*. Applied and Environmental Microbiology, 72(2), 1239–1247. 10.1128/AEM.72.2.1239-1247.2006.

