## Supplementary Material for "Structural models predict a significantly higher binding affinity between the NblA protein of cyanophage Ma-LMM01 and the phycocyanin of *Microcystis aeruginosa* NIES-298 compared to the host homolog"

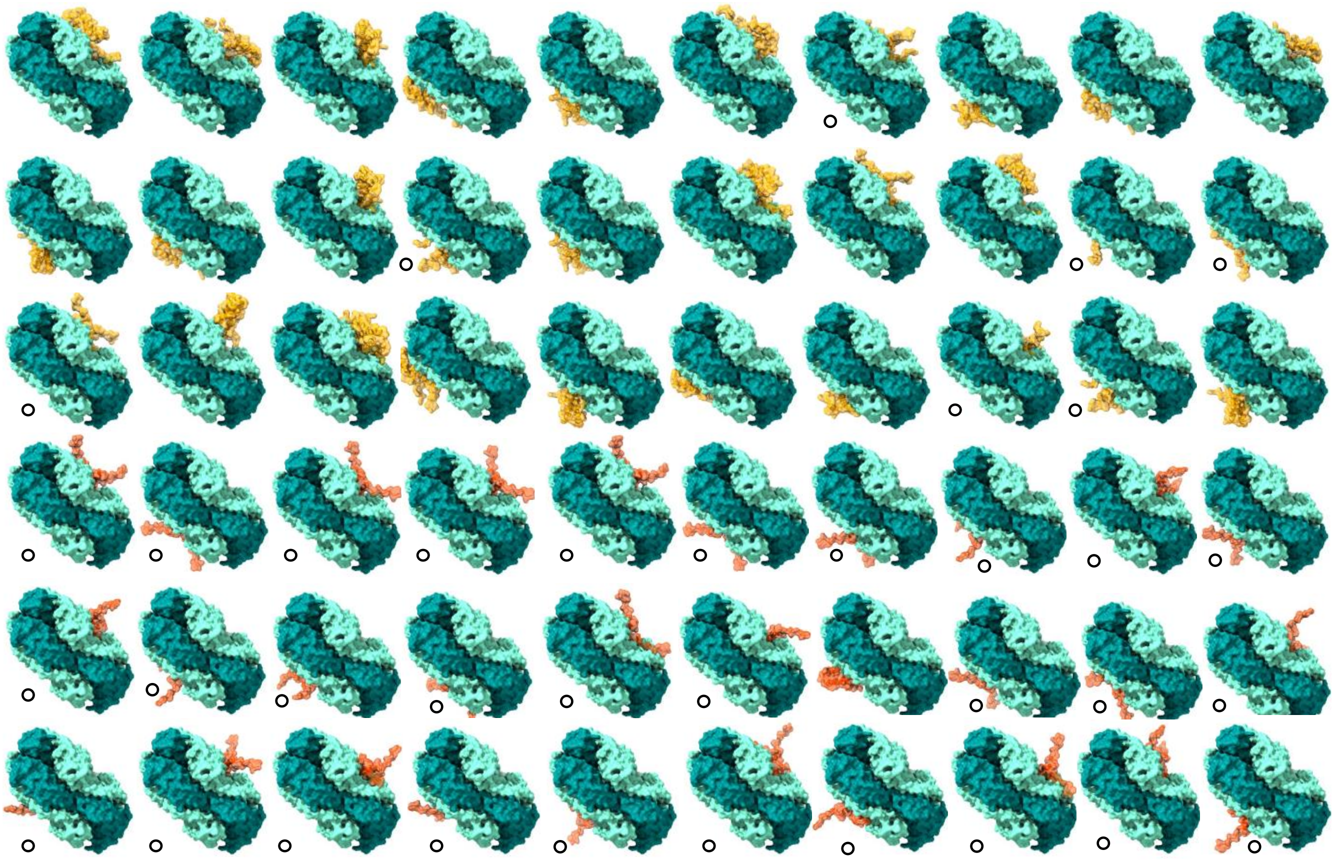

**Fig. S1**. Side views of the 30 top-ranked Ma-LMM01 non-bleaching A dimer (vNblA)- and *Microcystis aeruginosa* NIES-298 NblA dimer (hNblA)-*M. aeruginosa* NIES-298 phycocyanin $(\alpha\beta)_6$ hexamer (PC) Balanced ClusPro 2.0 docking models. The first three rows from top to bottom display the hNblA-PC complexes, while the last three show the vNblA-PC complexes. hNblA is colored in orange, vNblA in orange red; PC α- and β-subunits are colored in dark cyan and aquamarine, respectively. The docking models in which NblA is bound deep inside the PC groove are denoted with an open circle. Most hNblA models (23/30) were bound superficially (compared to vNblA) to PC, with a smaller region inside the PC groove. In contrast, virtually all vNblA models were bound deep inside the PC groove (29/30).

**Table S1**. Binding affinity prediction data of the *Microcystis aeruginosa* NIES-298 NblA-*M. aeruginosa* NIES-298 phycocyanin docking models. ICs, intermolecular contacts.

| Protein-protein complex | ΔG (kcal $mol^{-1}$) | $K_d$ (M) at 25 °C | ICs charged-charged | ICs charged-polar | ICs charged-nonpolar | ICs polar-polar | ICs polar-nonpolar | ICs nonpolar-nonpolar |
|---|---|---|---|---|---|---|---|---|
| model_000_00 | −14.9 | $1.20 \times 10^{-11}$ | 12 | 17 | 46 | 6 | 24 | 37 |
| model_000_01 | −15 | $1.00 \times 10^{-11}$ | 15 | 19 | 51 | 6 | 21 | 37 |
| model_000_02 | −16 | $1.90 \times 10^{-12}$ | 12 | 16 | 44 | 4 | 28 | 37 |
| model_000_03 | −12.6 | $6.00 \times 10^{-10}$ | 14 | 14 | 37 | 6 | 17 | 35 |
| model_000_04 | −16.1 | $1.60 \times 10^{-12}$ | 13 | 15 | 46 | 5 | 28 | 40 |
| model_000_05 | −13.4 | $1.40 \times 10^{-10}$ | 8 | 18 | 27 | 5 | 27 | 35 |
| model_000_06 | −13.1 | $2.60 \times 10^{-10}$ | 17 | 30 | 46 | 12 | 19 | 31 |
| model_000_07 | −16.1 | $1.70 \times 10^{-12}$ | 18 | 27 | 37 | 11 | 35 | 44 |
| model_000_08 | −17.2 | $2.50 \times 10^{-13}$ | 21 | 26 | 46 | 9 | 33 | 37 |
| model_000_09 | −18.4 | $2.90 \times 10^{-14}$ | 17 | 22 | 58 | 9 | 35 | 38 |
| model_000_10 | −24 | $2.50 \times 10^{-18}$ | 21 | 37 | 76 | 9 | 50 | 53 |
| model_000_11 | −16.2 | $1.30 \times 10^{-12}$ | 18 | 26 | 36 | 11 | 36 | 37 |
| model_000_12 | −17.4 | $1.90 \times 10^{-13}$ | 16 | 21 | 51 | 7 | 32 | 32 |
| model_000_13 | −14.9 | $1.20 \times 10^{-11}$ | 13 | 23 | 38 | 7 | 28 | 41 |
| model_000_14 | −15.4 | $5.40 \times 10^{-12}$ | 12 | 15 | 49 | 5 | 24 | 39 |
| model_000_15 | −14.3 | $3.40 \times 10^{-11}$ | 9 | 23 | 23 | 4 | 31 | 31 |
| model_000_16 | −16.4 | $9.10 \times 10^{-13}$ | 16 | 16 | 26 | 6 | 38 | 39 |
| model_000_17 | −18.6 | $2.10 \times 10^{-14}$ | 16 | 25 | 52 | 8 | 38 | 36 |
| model_000_18 | −16.8 | $5.10 \times 10^{-13}$ | 11 | 23 | 29 | 8 | 42 | 29 |
| model_000_19 | −13.3 | $1.70 \times 10^{-10}$ | 13 | 25 | 35 | 10 | 25 | 34 |
| model_000_20 | −11.9 | $1.80 \times 10^{-09}$ | 11 | 25 | 30 | 9 | 21 | 42 |
| model_000_21 | −15.5 | $4.20 \times 10^{-12}$ | 8 | 12 | 46 | 3 | 26 | 43 |
| model_000_22 | −15.8 | $2.60 \times 10^{-12}$ | 14 | 22 | 49 | 6 | 26 | 31 |
| model_000_23 | −11.9 | $1.80 \times 10^{-09}$ | 15 | 16 | 31 | 10 | 20 | 23 |
| model_000_24 | −17.7 | $1.10 \times 10^{-13}$ | 14 | 20 | 49 | 7 | 35 | 33 |
| model_000_25 | −21.4 | $2.20 \times 10^{-16}$ | 19 | 27 | 79 | 8 | 37 | 43 |
| model_000_26 | −16.5 | $7.60 \times 10^{-13}$ | 17 | 29 | 41 | 9 | 34 | 42 |
| model_000_27 | −18.3 | $3.90 \times 10^{-14}$ | 14 | 24 | 30 | 8 | 47 | 27 |
| model_000_28 | −11.8 | $2.10 \times 10^{-09}$ | 9 | 22 | 29 | 9 | 22 | 38 |
| model_000_29 | −15.8 | $2.60 \times 10^{-12}$ | 13 | 17 | 46 | 4 | 26 | 35 |

**Table S2**. Binding affinity prediction data of the Ma-LMM01 NblA-*Microcystis aeruginosa* NIES-298 phycocyanin docking models. ICs, intermolecular contacts.

| Protein-protein complex | ΔG (kcal mol$^{-1}$) | $K_d$ (M) at 25 °C | ICs charged-charged | ICs charged-polar | ICs charged-nonpolar | ICs polar-polar | ICs polar-nonpolar | ICs nonpolar-nonpolar |
|---|---|---|---|---|---|---|---|---|
| model_000_00 | −16.3 | $1.10 \times 10^{-12}$ | 21 | 23 | 48 | 4 | 25 | 34 |
| model_000_01 | −16.9 | $4.20 \times 10^{-13}$ | 20 | 25 | 48 | 4 | 28 | 35 |
| model_000_02 | −13.3 | $1.80 \times 10^{-10}$ | 21 | 20 | 36 | 3 | 16 | 32 |
| model_000_03 | −15.2 | $7.00 \times 10^{-12}$ | 14 | 20 | 42 | 1 | 23 | 26 |
| model_000_04 | −17 | $3.20 \times 10^{-13}$ | 18 | 25 | 51 | 5 | 29 | 40 |
| model_000_05 | −16.4 | $9.00 \times 10^{-13}$ | 18 | 22 | 43 | 3 | 28 | 31 |
| model_000_06 | −18.1 | $4.90 \times 10^{-14}$ | 23 | 24 | 51 | 2 | 29 | 35 |
| model_000_07 | −15.2 | $7.00 \times 10^{-12}$ | 22 | 20 | 36 | 4 | 25 | 31 |
| model_000_08 | −17.4 | $1.80 \times 10^{-13}$ | 21 | 24 | 39 | 2 | 32 | 31 |
| model_000_09 | −17 | $3.20 \times 10^{-13}$ | 22 | 24 | 45 | 4 | 29 | 36 |
| model_000_10 | −17.4 | $1.80 \times 10^{-13}$ | 18 | 25 | 52 | 5 | 30 | 37 |
| model_000_11 | −16.6 | $6.70 \times 10^{-13}$ | 17 | 25 | 50 | 4 | 27 | 33 |
| model_000_12 | −16.8 | $4.40 \times 10^{-13}$ | 25 | 28 | 54 | 3 | 22 | 41 |
| model_000_13 | −18.5 | $2.50 \times 10^{-14}$ | 25 | 22 | 47 | 8 | 37 | 33 |
| model_000_14 | −19 | $1.30 \times 10^{-14}$ | 24 | 15 | 49 | 3 | 34 | 33 |
| model_000_15 | −19.5 | $5.30 \times 10^{-15}$ | 23 | 19 | 59 | 5 | 34 | 33 |
| model_000_16 | −19.9 | $2.50 \times 10^{-15}$ | 26 | 21 | 68 | 9 | 34 | 55 |
| model_000_17 | −19.6 | $4.50 \times 10^{-15}$ | 23 | 28 | 66 | 8 | 34 | 31 |
| model_000_18 | −18.8 | $1.60 \times 10^{-14}$ | 24 | 22 | 48 | 8 | 38 | 36 |
| model_000_19 | −17.8 | $8.20 \times 10^{-14}$ | 23 | 19 | 50 | 10 | 35 | 32 |
| model_000_20 | −19.4 | $5.80 \times 10^{-15}$ | 31 | 21 | 60 | 12 | 36 | 33 |
| model_000_21 | −16.1 | $1.50 \times 10^{-12}$ | 15 | 21 | 47 | 5 | 28 | 22 |
| model_000_22 | −15.5 | $4.10 \times 10^{-12}$ | 26 | 30 | 51 | 12 | 25 | 43 |
| model_000_23 | −18.9 | $1.40 \times 10^{-14}$ | 24 | 19 | 52 | 6 | 35 | 36 |
| model_000_24 | −17.9 | $7.00 \times 10^{-14}$ | 28 | 39 | 59 | 13 | 32 | 43 |
| model_000_25 | −19.9 | $2.70 \times 10^{-15}$ | 29 | 36 | 67 | 12 | 36 | 39 |
| model_000_26 | −20.3 | $1.30 \times 10^{-15}$ | 38 | 37 | 72 | 11 | 31 | 38 |
| model_000_27 | −20.4 | $1.20 \times 10^{-15}$ | 19 | 26 | 63 | 4 | 37 | 36 |
| model_000_28 | −18.4 | $3.10 \times 10^{-14}$ | 23 | 22 | 55 | 7 | 33 | 30 |
| model_000_29 | −16.2 | $1.40 \times 10^{-12}$ | 29 | 36 | 52 | 12 | 26 | 40 |
